# A chromosome-level genome resource for the lethal tapeworm *Sparganum proliferum*, with candidate genomic safe harbours for functional genetics

**DOI:** 10.1101/2025.09.02.673767

**Authors:** Atsushi Hikima, Simo Sun, Tetsuya Okino, Kensei Kinoshita, Akemi Yoshida, Yasunobu Maeda, Kenji Ishiwata, Hirotaka Kanuka, Taisei Kikuchi

## Abstract

*Sparganum proliferum* is an enigmatic and highly proliferative cestode known for its ability to form invasive larval infections in mammalian hosts. Here, we present the first chromosome-level genome assembly for a member of the order Diphyllobothriidea, generated using a hybrid sequencing approach that integrates Oxford Nanopore/PacBio long-read sequencing, Illumina short-read sequencing, and Hi-C scaffolding. The final genome assembly spanned approximately 681 megabases (Mb) across nine chromosomes. BUSCO analysis revealed 81.9% completeness, whereas repeat annotation identified 55.8% of the genome as repetitive elements. Gene annotation uncovered approximately 29,000 protein-coding genes including ∼6200 transposon associated genes, highlighting the complex genomic landscape underlying parasitic lifestyles. Synteny analysis with other cestode linages including *Echinococcus* and *Hymenolepis* provided insights into the structural organisation and evolutionary trajectory of *S. proliferum*. In addition, we identified candidate genomic safe harbour (GSH) loci and promoters from housekeeping genes, offering potential for stable transgene integration. This high-quality genome serves as a critical resource for studying parasite evolution, host adaptation mechanisms, and the molecular basis of invasive proliferation in cestodes.

## Introduction

Tapeworms (class Cestoda) represent a diverse and evolutionarily specialised group of endoparasitic flatworms with complex lifestyles that often involve multiple intermediate hosts ^1^. Their parasitic lifestyle has driven significant genomic and biological adaptations, particularly in the context of host interaction, immune evasion, and nutrient acquisition. Among cestodes, *Sparganum proliferum* stands out due to its capacity to undergo continuous larval proliferation in infected mammalian hosts, leading to severe sparganosis ^2^. Unlike most tapeworms that develop into adult forms in the definitive hosts, *S. proliferum* is characterised by continuous and invasive asexual proliferation during its larval stage. Documented cases of *S. proliferum* are rare; however, infection in humans is fatal in most cases. The infection routes and the mechanisms underlying asexual multiplication and its pathogenesis remain unclear.

Despite the clinical significance and evolutionary distinctiveness of *S. proliferum*, our understanding of its biology at the molecular level remains limited. The genome of *S. proliferum* was previously sequenced, revealing its close phylogenetic relationship with non-proliferative *Spirometra* species and identifying expanded gene families that may play a role in parasitism and host interaction ^3^. However, due to the genome’s repetitive nature, the assembly was fragmented, which restricts deeper insight into chromosomal organisation, gene regulation, and the genetic basis of its proliferative capabilities.

In this study, we leveraged a combination of Oxford Nanopore, PacBio HiFi, Illumina short reads, and Hi-C sequencing to produce a chromosome-level genome assembly of *S. proliferum*. The resulting assembly is, to our knowledge, the most complete genome for any member of the order Diphyllobothriidea. This high-quality genomic resource enabled the annotation of over 29,000 genes and a comprehensive analysis of repeat elements, chromosomal synteny with related species, and gene family evolution. Furthermore, we identified candidate genomic safe harbour (GSH) regions— transcriptionally active intergenic loci that are amenable to stable transgene insertion. These loci and associated promoter regions will serve as foundational tools for future genetic manipulation and functional studies in cestodes.

## Results and Discussion

### Genome Assembly and Structural Organization

We produced a chromosome-level genome assembly of *S. proliferum* using Oxford Nanopore/PacBio long-read sequencing for de novo assembly, Illumina sequencing for base-error correction, and Hi-C scaffolding. Details of the raw sequence data are provided in Table S1. The final 681 Mb genome assembly comprised 32 scaffolds, with 99.8% of the total assembled bases contained within the top 10 scaffolds. Hi-C chromatin interaction data revealed strong intra-chromosomal interactions along the diagonals of the nine main scaffolds (Fig. 1). In contrast, the 10th largest scaffold, measuring approximately 16.4 Mb, exhibited weaker Hi-C interaction signals at both intra- and inter-chromosomal levels. This indicates that this scaffold may have a distinct nature from the other nine pseudo-chromosomal scaffolds. Karyotype analysis verified that *S. proliferum* has a chromosome complement of 2n = 18, providing a cytogenetic basis for genome organization (Fig. 2).

**Fig. 1.**
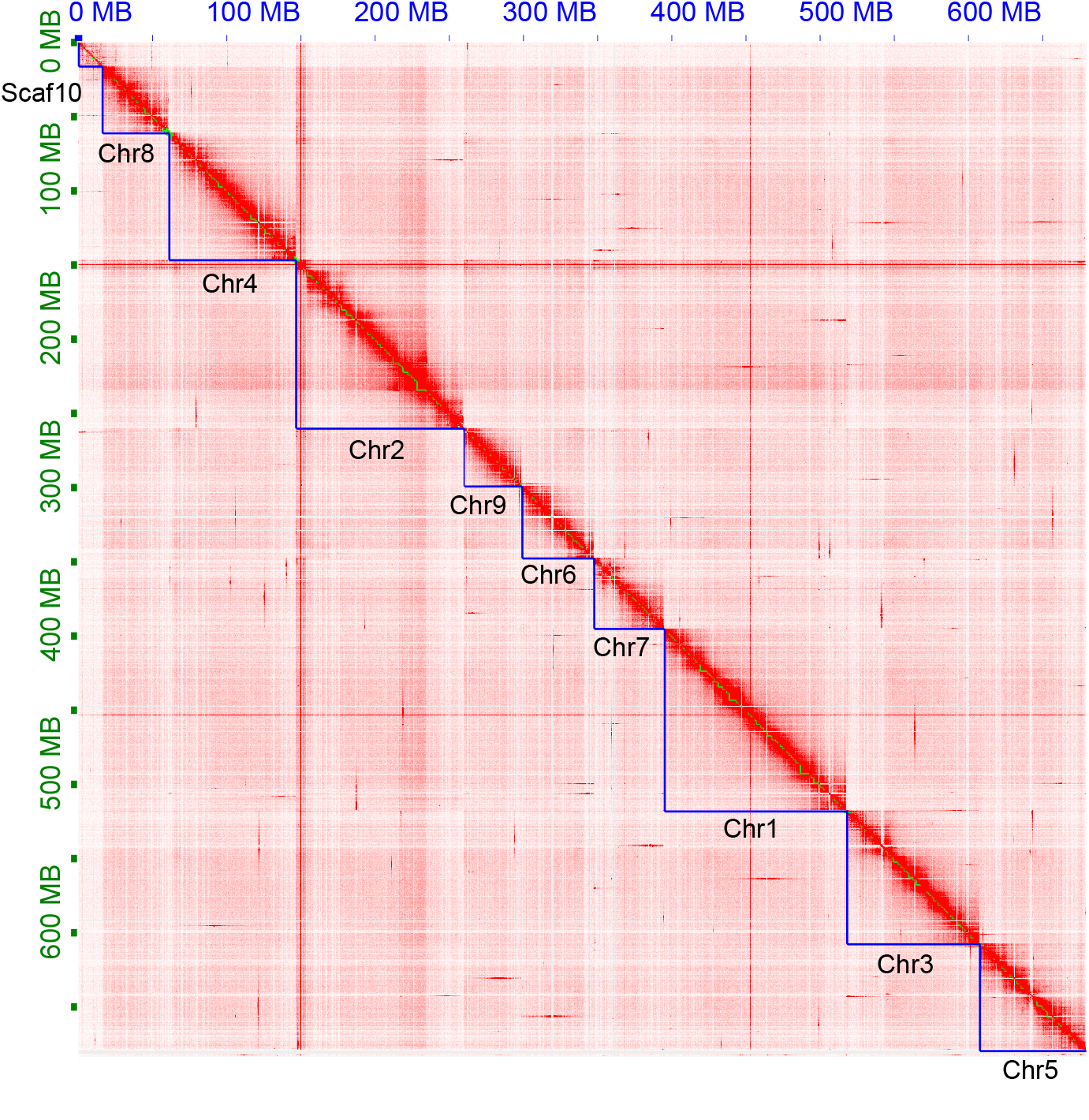
Hi-C contact map of the *Sparganum proliferum* genome assembly. Stronger signal intensity (darker color) represents higher contact frequency between genomic loci.

**Fig. 2.**
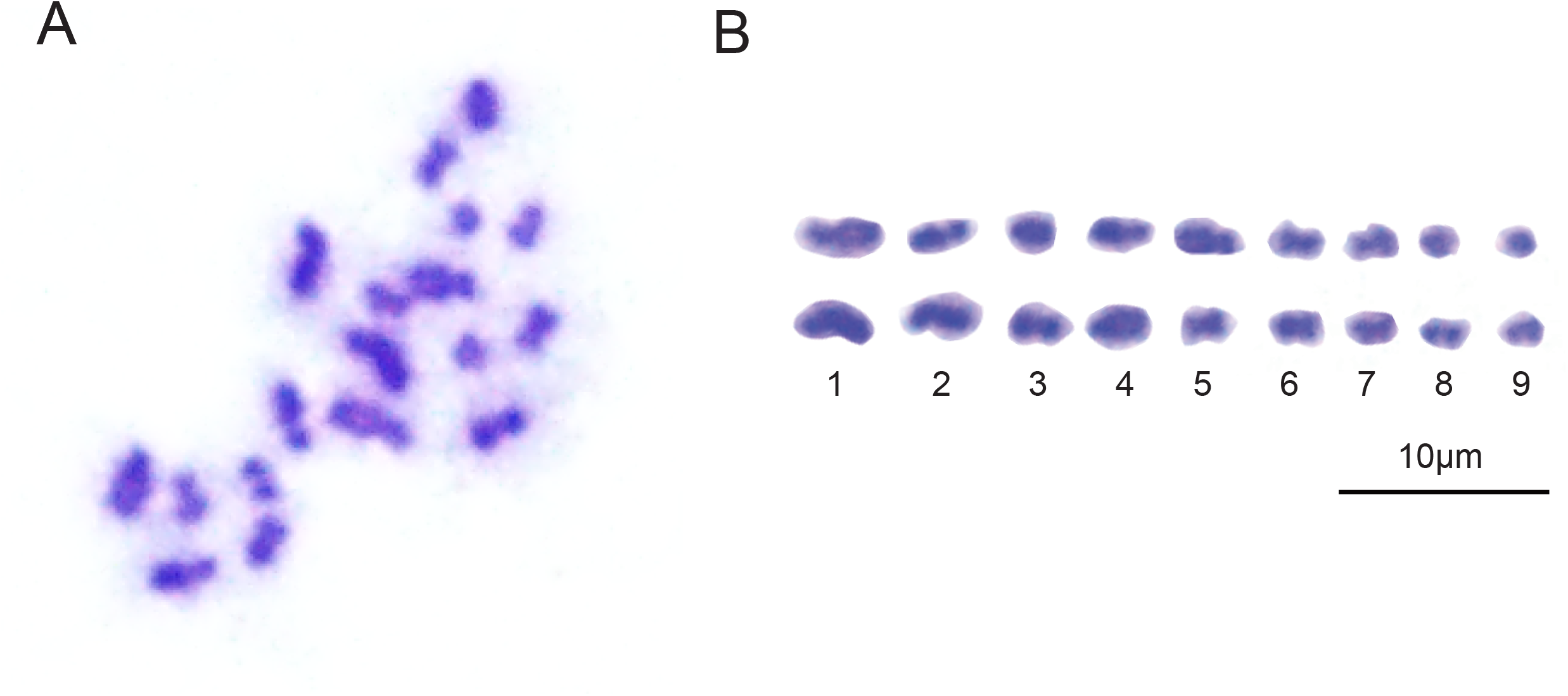
Mitotic chromosome spreads of *Sparganum proliferum*. (A) Representative metaphase chromosome spread of *S. proliferum* prepared from plerocercoids using colchicine treatment, stained with Giemsa. Chromosomes appear as condensed rod-like structures. Nine chromosome pairs (n = 9) are visible. (B) Karyogram showing the arranged and numbered chromosomes based on relative size and morphology. Scale bar = 10 μm.

### Assembly Quality and Completeness

The BUSCO (Benchmarking Universal Single-Copy Orthologs) analysis demonstrated 81.9% completeness against the metazoan database, a value that exceeds the previous short-read assembly (v2.2) and those reported for most other cestode genomes (Table 1). For example, the chromosome-scale genome of *Hymenolepis microstoma* yielded 82.4% BUSCO assembly completeness ^4,5^, while *Echinococcus granulosus achieved* 82.8% ^6^. These values are representative of a broader trend across cestode genomics, where BUSCO completeness tends to be lower than in many other metazoan groups, likely due to high repeat content, genome plasticity, and annotation challenges inherent to parasitic flatworms ^6^. These comparisons underscore the relative success of the *S. proliferum* assembly in capturing a more complete set of conserved genes, particularly given the inherent difficulties in assembling and annotating cestode genomes.

**Table 1.** Statistics of the new chromosome-level genome assembly (v4) of *Sparganum proliferum* compared with the previous assembly and other flatworm species.

|  | *Sparganum*<br>*proliferum* (v4) | *Sparganum*<br>*proliferum*<br>(v2.2) | *Spirometra*<br>*mansoni*<br>(v2.0) | *Hymenolepis*<br>*microstoma*<br>(WBPS19) | *Echinococcus*<br>*granulosus*<br>(WBPS19) | *Schmidtea*<br>*mediterranea*<br>(WBPS19) |
| --- | --- | --- | --- | --- | --- | --- |
| Assembly size<br>(Mb) | 681 | 653 | 796 | 169 | 172 | 840 |
| Num. scaffolds | 32 | 7,388 | 5,723 | 7 | 31 | 656 |
| Largest scaff<br>(kb) | 123,237 | 8,099 | 5,490 | 43,032 | 38,513 | 339,756 |
| N50 (kb) | 85,402 | 1,242 | 821 | 25,815 | 18,675 | 270,168 |
| N90 (kb) | 44,839 | 110 | 167 | 17,547 | 12,341 | 55,279 |
| Gaps (kb) | 1,070 | 51,020 | 77,788 | 6,487 | 4,025 | 17 |
| BUSCO<br>(assembly)<br>completeness<br>(Eukaryota<br>dataset) | 81.9 | 79.2 | 79.6 | 82.4 | 82.8 | 80.0 |
| Num. genes<br>(Coding genes<br>/Transposable<br>element) | 29,231<br>(22,915/6,316) | 23,043<br>(18,919/4,124) | 28,290<br>(22,162/6,128) | 11429<br>(-/-) | 9985<br>(-/-) | 33445<br>(-/-) |
| Average<br>protein length | 448 | 401 | 407 | 576 | 516 | 482 |

### Gene Annotation and Repeats

A total of 29,231 genes were predicted, marking an increase of approximately 6,200 genes over the earlier assembly (Table 1). Of these, 6,316 genes were identified as encoding transposable element proteins, while the remaining 22,915 were annotated as standard protein-coding genes. Repeat elements comprised about 380 Mb, making up 55.77% of the genome (Table 2).

**Table 2.**
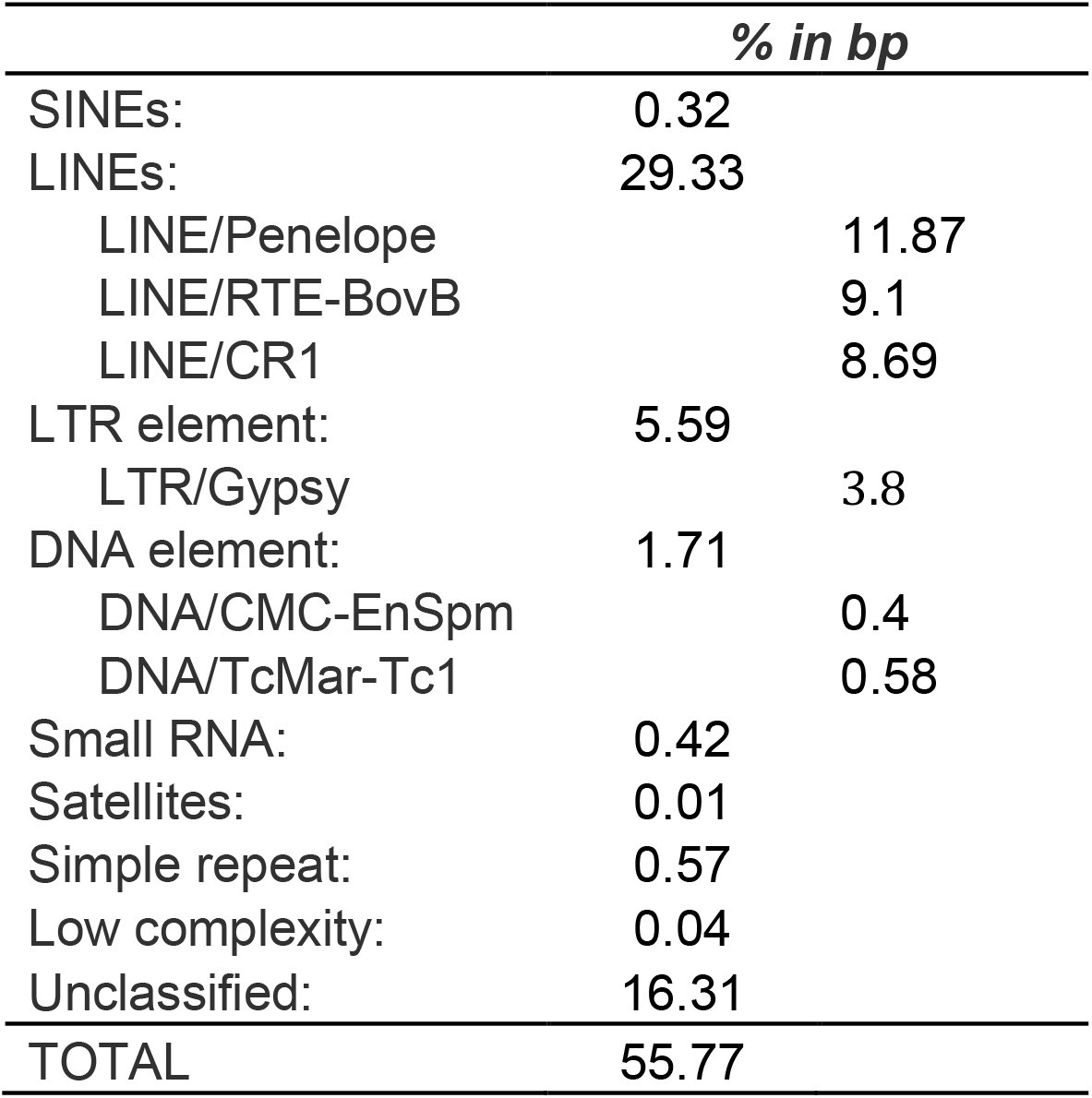
Repeat content composition of the *Sparganum proliferum* v4 genome assembly.

|  | % *in bp* |  |
| --- | --- | --- |
| SINEs: | 0.32 |  |
| LINEs: | 29.33 |  |
| LINE/Penelope |  | 11.87 |
| LINE/RTE-BovB |  | 9.1 |
| LINE/CR1 |  | 8.69 |
| LTR element: | 5.59 |  |
| LTR/Gypsy |  | 3.8 |
| DNA element: | 1.71 |  |
| DNA/CMC-EnSpm |  | 0.4 |
| DNA/TcMar-Tc1 |  | 0.58 |
| Small RNA: | 0.42 |  |
| Satellites: | 0.01 |  |
| Simple repeat: | 0.57 |  |
| Low complexity: | 0.04 |  |
| Unclassified: | 16.31 |  |
| TOTAL | 55.77 |  |

### Synteny and Comparative Genomics

We investigated chromosomal synteny between *S. proliferum* and three other flatworm species with chromosome-level genome assemblies: *H. microstoma, E. granulosus*, and the non-parasitic planarian *Schmidtea mediterranea*. There was minimal conserved synteny between *S. proliferum* and *S. mediterranea*, reflecting their deep evolutionary divergence (Fig. 3). In contrast, *S. proliferum* chromosomes showed a well conserved synteny with the parasitic cestodes, especially *E. granulosus*, where the chromosomes demonstrated a distinct one-to-one correspondence. These findings support the notion of lineage-specific chromosomal evolution and rearrangements associated with lifestyle and host adaptation.

**Fig. 3.**
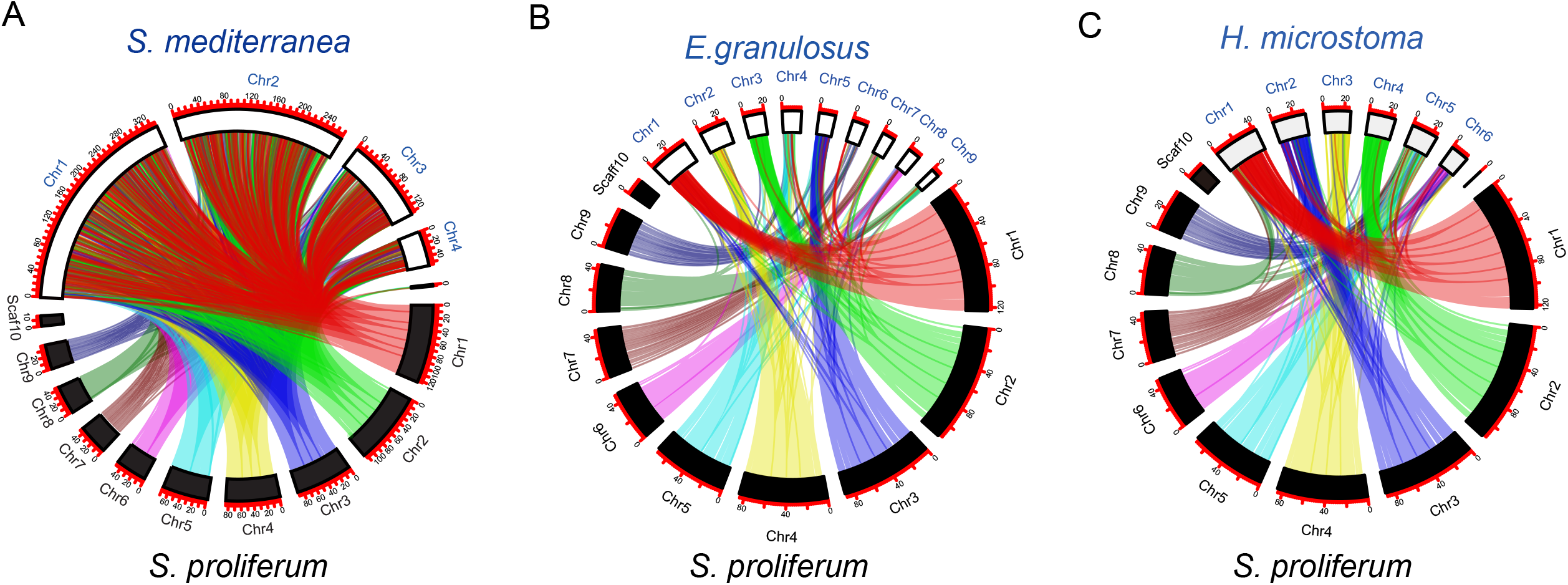
Synteny comparison of *Sparganum proliferum* with other flatworm genomes. Circos plots showing macrosyntenic relationships between the chromosome-level genome assembly of S. proliferum (black labels) and three other flatworm species. A) Comparison with the free-living planarian *Schmidtea mediterranea* reveals extensive genome rearrangements, reflecting deep divergence between parasitic and non-parasitic flatworms. B) Comparison with *Hymenolepis microstoma* shows higher synteny conservation with fewer large-scale rearrangements. C) Comparison with *Echinococcus granulosus* also highlights significant syntenic conservation with species-specific differences. Colored ribbons represent orthologous gene pairs. Black and white boxes denote chromosomes (or scaffolds), with the scale in megabases (Mb) indicated along the outer edges.

### Genomic Safe Harbours and Promoter Discovery

To facilitate future genome editing and transgenesis, we sought to identify genomic safe harbours (GSHs)—intergenic loci capable of accommodating exogenous DNA without disrupting native gene function. We defined GSHs as transcriptionally active regions located at least 2 kb from the nearest gene with convergent orientation and containing PAM sequences (NGG), which are essential for CRISPR-Cas9 targeting. Those are located in Chromosome 2 and 4 (Table 3, Table S2-S4). We also screened upstream regions of housekeeping genes to identify candidate promoters suitable for driving stable transgene expression (Table S5). These resources represent a significant advance toward the development of functional genomic tools in cestodes, enabling controlled gene knock-ins and expression studies.

**Table 3.** Genomic locations and characteristics of genome safe harbor (GSH) sites for CRISPR-cas9 targeting in the *Sparganum proliferum* v4 assembly.

| GSH name | chromosome | location | length (bp) | number of CRISPR-cas9 target sequences |
| --- | --- | --- | --- | --- |
| GSH1 | SPRV4.Chr02 | 34566861-34568808 | 1,947 | 10 (Table S2) |
| GSH2 | SPRV4.Chr02 | 46400640-46405293 | 4,653 | 9 (Table S3) |
| GSH3 | SPRV4.Chr04 | 22169935-22173224 | 3,289 | 7 (Table S4) |

## Conclusion

The availability of a chromosome-level genome assembly for *S. proliferum*, as the first high quality genome reference of Diphyllobothriidea, opens new avenues for exploring the genetic basis of its pathogenicity and its unusual capacity for larval proliferation. The identification of syntenic regions and expanded gene families involved in host-parasite interaction may yield insights into the molecular mechanisms of immune evasion and tissue invasion. Additionally, the GSH and promoter candidates provide a crucial foundation for implementing CRISPR-based technologies and transgenesis protocols in tapeworms—tools that are currently lacking in most parasitic flatworms. These advances collectively lay the groundwork for functional studies that can dissect gene function and regulatory networks in *S. proliferum* and related species.

## Methods

### Ethics statement

All procedures involving vertebrate animals were approved by the Institutional Animal Care and Use Committee of The University of Tokyo (protocol #A2023FS004) and complied with national regulations and institutional guidelines. Animal numbers were minimized and procedures refined in accordance with the 3Rs. Reporting follows the ARRIVE 2.0 guidelines; key husbandry, monitoring, and humane endpoint details are provided in the Methods.

### Parasite maintenance and DNA extraction

*S. proliferum* strain Venezuela used in this study has been maintained by serial passages using BALB/c mice via intraperitoneal injections of the plerocercoids at the Laboratory of Parasite System Biology, the University of Tokyo as described in De Noya et al ^7^. *S. proliferum* larvae were collected from the abdominal cavity of infected mice and washed thoroughly with 1x PBS. The worms were incubated at 60°C for 1 h with a solution comprising 100 µL Proteinase K (Wako), 100 µL of 1M DTT (Wako), and 4 mL G2 buffer (Qiagen). Then, 70 µL RNase (Invitrogen) was added and incubated at 37°C for 30 min. The lysate was treated with phenol/chloroform extraction followed by purification using a Genomic-tip 20/G according to the manufacturer’s protocol (Qiagen). For long-read sequencing short DNA fragment removal treatment was applied using the SRE KIT (PacBio). The DNA quality and concentration were assessed using TapeStation (Agilent) and a Qubit 2.0 Fluorometer (Life Technologies, CA, USA).

### DNA sequencing and Hi-C analysis

The genome sequencing and assembly process utilised a multiplatform approach. Oxford Nanopore sequencing produced approximately 11.1 Gb of long-read data, with an N50 read length of 13.2 kb, while PacBio HiFi sequencing produced yielding 21.6 Gb with an N50 read length of 10.3 kb. Illumina short-read data were obtained from a previous study^3^ with 72.0 Gb of paired-end 150 base-pair (bp) reads. Hi-C sequencing was performed using an ARIMA kit to capture chromatin conformation (ARIMA), producing 25.8 Gb of data. De novo genome assembly was conducted using NextDenovo ^8^, followed by three rounds of base error correction using Pilon ^9^ and the Illumina short reads. Hi-C data were processed using Juicer ^10^ and 3D-DNA ^11^ pipelines, yielding a chromosome-level assembly.

### Chromosome observation

Karyotype examination was conducted using Giemsa staining of worms treated with colchicine-Hank’s solution (Wako) at 37°C for 4 h. Manual enumeration of chromosomes was performed on the captured images, which revealed that the majority of nuclei contained 18 chromosomes (Fig. 2). Based on these observations, we concluded that the karyotype of *S. proliferum* is 2n = 18. This finding aligns with those of *Spirometra mansoni*, a closely related species that has been confirmed to possess nine haploid chromosomes through the observation of triploid specimens ^12^.

### Genome annotation

AUGUSTUS v. 3.2.3^13^ was used to predict the protein-coding genes. RNA-seq reads were used to generate intron hints using Hisat2 v2.2.1^14^ and bam2hints included in the AUGUSTUS package. AUGUSTUS was executed using intron hints and species parameters trained through manually curated training sets^3^. The predicted gene set was functionally annotated using eggNOG-mapper v2.1.12^15^ to assign GO terms and protein families. TransposonPSI (https://transposonpsi.sourceforge.net) was employed to detect transposable element proteins. Repeat analysis was conducted using RepeatModeler (https://github.com/Dfam-consortium/RepeatModeler) and RepeatMasker (http://www.repeatmasker.org/) software.

### Gene Orthology and Synteny

OrthoFinder v.2.5.4^16^ was used to obtain one-to-one single copy orthologous of *S. proliferum* and the three other platyhelminth species including *H. microstoma, E. granulosus*, and *S. mediterranea* whose data was retrieved from WormBase Parasite WBPS19 (https://parasite.wormbase.org/index.html). TBtools^17^ was used to visualise the positions of these orthologous genes on the chromosomes of each species.

### Identification of promoters and Genomic safe harbour

Previously published RNA-seq data were used for transcriptional profiling of *S. proliferum* larvae ^3^. The dataset included six plerocercoid samples, comprising three medusa-form larvae (vigorous budding phenotype) and three wasabi-form larvae (clotted, non-proliferative phenotype). RNA-seq reads were mapped to the *S. proliferum* genome assembly (v4.0) using HISAT2 v2.2.1^14^ with default parameters and FPKM (Fragments Per Kilobase of transcript per Million mapped reads) values for all predicted genes were calculated using Cufflinks v2.2.1^18^.

To identify transcriptionally active regions, a sliding window of 10 consecutive genes was applied across the genome. A region was defined as transcriptionally active if the average FPKM of the 10 genes exceeded 10 in all six RNA-seq samples. In addition, the total intergenic distance across the 10-gene window was required to be less than 10 kb, ensuring the region is both transcriptionally active and gene-dense.

Candidate genomic safe harbours (GSHs) were defined as intergenic regions that met the following criteria:

1. Located within transcriptionally active regions as defined above,
2. At least 2 kb from the nearest annotated gene,
3. Flanked by convergently oriented genes, and
4. Containing at least one CRISPR/Cas9 PAM sequence (NGG).

The Safe Harbor Identification Program (SHIP) ^19^ and custom scripts were used to extract intergenic coordinates from the genome annotation GFF file and intersect them with gene expression windows. PAM motifs (identified via regular expression matching of “NGG” in the reference sequence) were identified using ChopChop ^20^.

Promoter regions (1–2 kb upstream) of housekeeping genes were extracted for potential use in transgene expression. Housekeeping genes were selected based on eggNOG-mapper annotations and BUSCO single-copy orthologs. Genes showing consistently high FPKM values across all six RNA-seq samples were prioritized, including well-conserved genes such as ef-1a and tubulin.

## Data Availability

All sequencing reads, including those from Oxford Nanopore, Illumina, and Hi-C technologies, have been deposited in DRA under accession number DRR684723-DRR684725 (Table S1). The assembled genome and associated annotations are available at DDBJ/ENA/GenBank under BioProject accession numbers PRJEB35374. RNA-seq data used for gene annotation are accessible under BioProject accession number PRJDB8966.

## Acknowledgments

We extend our gratitude to the members of the Parasite Systems Biology Lab in UTokyo for their useful advice and continuous support. This research was supported by the Japan Society for the Promotion of Science (JSPS) KAKENHI Grant Numbers 19H03212 and 25H01307, and JST CREST Grant Numbers JPMJCR18S7 and JPMJCR23B1.

## Author Contributions

TK and HK were responsible for conceptualisation and project administration. AH, SS, KI and AY conducted sample preparation, genome sequencing and assembly, whereas AH and YM performed genome annotation and comparative analyses. TO performed cytological analysis. TK and AH drafted the manuscript, with all authors contributing to the revisions and final approval.

